# Endogenous viral elements trace the ancient origins and early evolution of the *Caulimoviridae*

**DOI:** 10.1101/2025.07.25.666790

**Authors:** Héléna Vassilieff, Saad Serfraz, Nathalie Choisne, Andrew D.W. Geering, Pierre Lefeuvre, Pierre-Yves Teycheney, Florian Maumus

**Affiliations:** Université Paris-Saclay, INRAE, BioinfOmics, URGI, 78026, Versailles, France; CABB, University of Agriculture, Faisalabad, 38000, Pakistan; QAAFI, The University of Queensland, Brisbane, QLD 4072, Australia; CIRAD, UMR PVBMT, F-97410 Saint-Pierre de La Réunion, France; CIRAD, UMR PVBMT, Department of Plant Protection, College of Agriculture, Can Tho University, Can Tho City, Vietnam

## Abstract

Endogenous viral elements (EVEs) are viral sequences integrated into the genomes of host organisms, analogous to molecular fossils. The majority of characterised EVEs in plants are derived from the *Caulimoviridae*, the only family of dsDNA viruses infecting this kingdom. Endogenous caulimovirids (ECVs) occur across taxonomically diverse vascular plant species and represent a significant resource for studying host-virus coevolution, host range dynamics, and the evolution of viral genomes over deep timescales.

Previous evolutionary studies based on ECVs proposed either cospeciation or host-switching as the main drivers of *Caulimoviridae* evolution, but were limited by sparse genomic data from basal plant lineages. Using 93 plant genomes spanning all embryophyte lineages, including ferns and lycophytes, we identified 47,135 ECVs in 75 genomes. These were grouped into 71 operational taxonomic units (OTUs), including 35 novel ones, revealing unexpected *Caulimoviridae* diversity in vascular plants and a new clade restricted to gymnosperms. Phylogenetic comparisons with host plant taxonomy support a macroevolutionary scenario in which cospeciation with tracheophytes drove the diversification of *Caulimoviridae*. Our findings position *Caulimoviridae* and ECVs as a reference system for paleovirology, offering unprecedented insights into how plant viruses and their hosts coevolved, diversified, and sometimes went extinct.

## Introduction

The evolution of land plants has been significantly shaped by selective pressures imposed by pathogens, including viruses. Although numerous plant virus families have been identified ^1,2^, their origins and patterns of macroevolution remain poorly understood. Metagenomic surveys have revealed a wealth of previously undetected viruses and firmly established viruses as integral components of ecosystems, emphasizing the vast yet largely undocumented plant virosphere ^3^. However, viral metagenomics generally provides only a temporal snapshot of viral communities. In contrast, viral sequences integrated into host genomes, known as endogenous viral elements (EVEs), provide a unique, untargeted record of historical viral infections. Mining host genomes for EVEs has proven highly informative, as these molecular fossils allow the reconstruction of past viral diversity and host-virus interactions across evolutionary timescales, providing invaluable insights into the macroevolution of viruses ^4^.

In plants, most characterized EVEs are derived from the *Caulimoviridae*, the only family of dsDNA viruses in this kingdom, often referred to as plant pararetroviruses ^5^. Caulimovirids possess 7.1 - 9.8 kbp, circular, non-covalently closed double-stranded DNA genomes that contain a core suite of genes including a 30K movement protein (MP), a capsid protein (CP), a minor virion-associated protein (VAP) and a polymerase polyprotein (Pol) containing aspartic protease (AP), reverse transcriptase (RT), and ribonuclease H1 (RH) domains ^6^. They are currently classified into eleven genera based on virion morphology, genome organization, and transmission mode. Species are distinguished by RT-RH1 sequence divergence, with an 80% nucleotide identity threshold. Viruses in some genera encode auxiliary proteins, such as the aphid transmission factor in the genera *Caulimovirus* and *Soymovirus*, which allow these viruses to occupy specific ecological niches ^6^.

*Caulimoviridae* belongs to the order *Ortervirales*, along with *Retroviridae*, *Belpaoviridae* (Bel/Pao elements), *Metaviridae* (Ty3/Gypsy elements), and *Pseudoviridae* (Ty1/Copia elements) ^7^. All share a replication cycle alternating between dsDNA and ssRNA via reverse transcription. *Caulimoviridae* differ from other *Ortervirales* by lacking an integrase-mediated proviral stage. Nevertheless, their sequences are widely integrated into plant genomes, likely due to incorporation as filler DNA during the repair of double-strand DNA breaks in host chromosomes ^5^.

Since the first endogenous caulimovirids (ECVs) were reported in the late 1990s ^8–10^, the rapid advancement of plant genome sequencing has enabled their widespread detection using bioinformatics analysis. While only a few ECVs are replication-competent and capable of reactivation ^11–13^, dozens of complete or partial viral genomes have been assembled from ECVs without known episomal forms. These genomes encode putative replication, structural, and movement proteins typical of caulimovirids. Although phylogenetic analyses place them within the family, many stand outside the 108 species and eleven genera currently recognized by the International Committee for Taxonomy of Viruses (ICTV) ^6,14–19^. Most such viral genomes were assembled from ECVs belonging to the same family by aligning multiple, highly similar copies that share greater than 90% nucleotide identity in the region encoding the RT-RH1 domain.

Previous studies have shown that ECVs extend across all major euphyllophyte lineages (ferns, gymnosperms, and angiosperms), far exceeding the host range of known episomal caulimovirids ^5^. Mushegian and Elena (2015) ^15^ were the first to identify caulimovirid MP homologs in the genomes of gymnosperms and ferns. This host distribution was later confirmed and extended by Gong & Han (2018) ^17^ and Diop *et al.* (2018) ^16^ ; the latter study also reported the presence of a caulimovirid RT transcript in the lycophyte *Lycopodium annotinum*. However, ECV studies were constrained by a poor representation of genomic and transcriptomic data from basal plants. The surge in high-throughput sequencing, particularly of genomes from previously underrepresented taxa such as lycophytes, now enables a more comprehensive exploration of ECVs across tracheophytes. Automated bioinformatic pipelines for ECV detection and annotation ^20^ further facilitate host-range assessment and refinement of evolutionary hypotheses.

Here, we investigated ECV diversity in 93 plant genome assemblies and transcriptomes spanning basal and representative tracheophytes. Thousands of novel ECVs were identified, extending the *Caulimoviridae* host range to all tracheophyte divisions. By integrating these with prior studies ^14,16–19,21,22^, we propose a macroevolutionary model in which long-term host–virus cospeciation was punctuated by host and virus extinction events. Our findings position *Caulimoviridae* and their endogenous counterparts as a central model for investigating the evolution of plant viruses.

## Results

### Extended diversity and host range of the *Caulimoviridae*

We screened 73 plant genomes (Figure 1; Supp. Table 1) for the presence of endogenous caulimovirid RT domains (ECRTs) using the Caulifinder pipeline branch B described by Vassilieff *et al.* (2022) ^20^. The dataset included seven bryophytes and 66 tracheophytes comprising nine lycophytes, 17 ferns, 27 gymnosperms, three basal angiosperms (ANA-grade), and ten mesangiosperms (one species of *Chloranthaceae* and nine Magnoliids). This screening yielded 20,267 amino acid ECRT sequences (aa-ECRTs) across 55 plant species (Figure 1). No aa-ECRTs were detected in any of the seven bryophyte genomes analysed, nor in six lycophytes and five ferns (Figure 1). In contrast, aa-ECRTs were detected in the genomes of three lycophytes, twelve ferns, and all gymnosperm, basal angiosperm, *Chloranthaceae*, and Magnoliid species examined (Figure 1). The number of aa-ECRTs per genome varied greatly, even within the same plant division, particularly in gymnosperms, where the number of aa-ECRTs ranged from three in *Metasequoia glyptostroboides* to 3,352 in *Pinus tabuliformis*. A positive correlation was observed between the number of aa-ECRTs and host genome size (p-value = 9e-12). However, genome size alone accounted for only half of the variation (R² = 0.5; Supp. Figure 1).

**Figure 1:**
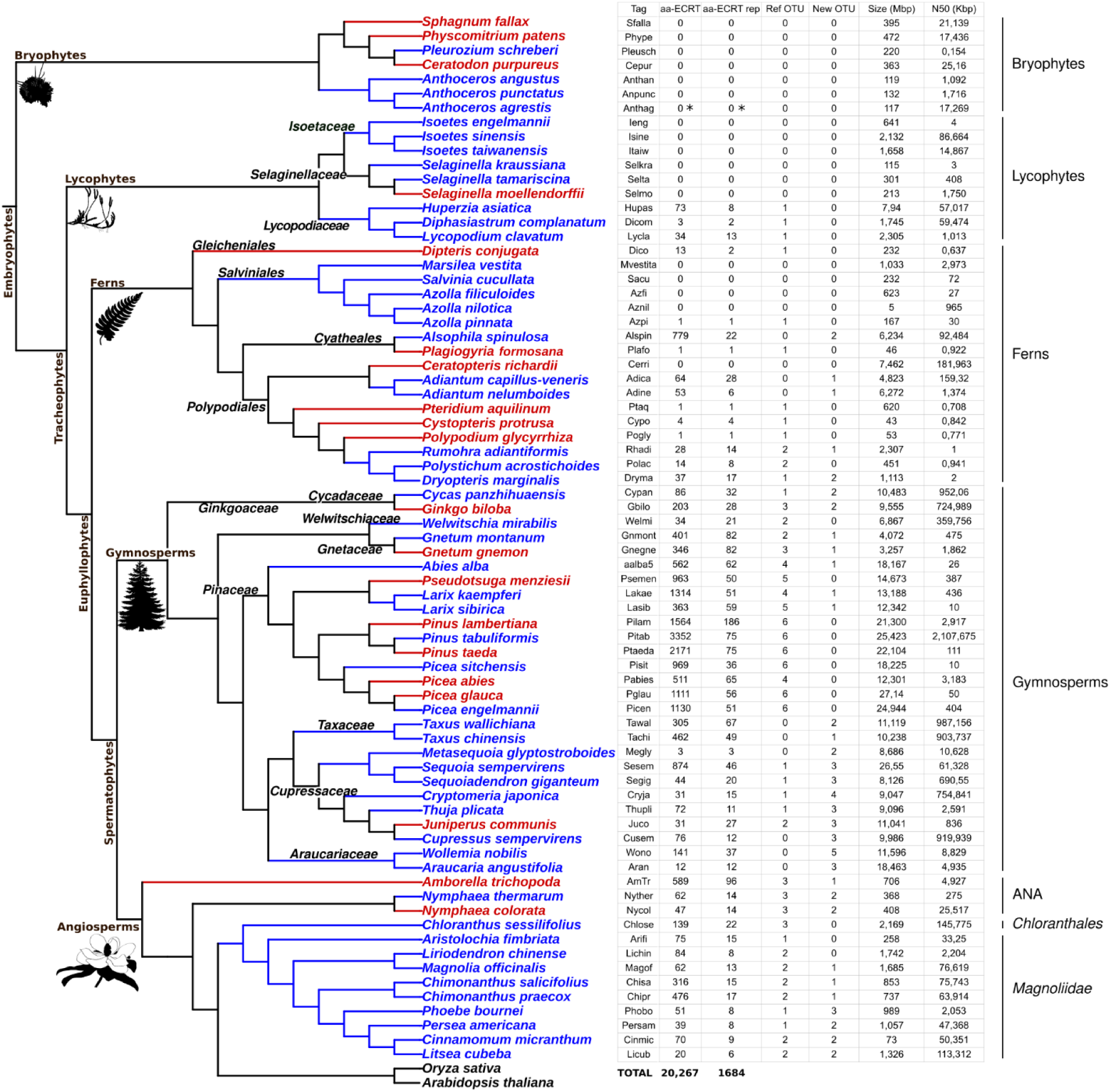
Distribution of aa-ECRTs, aa-ECRTs rep, and OTUs across plant genomes. The cladogram on the left illustrates the phylogenetic relationships among the 73 plant species analyzed. Species previously examined for ECVs detection are highlighted in red ^14–17,19,21^, while species investigated for the first time are highlighted in blue. Major plant lineages are annotated, including bryophytes, lycophytes, ferns, gymnosperms, and angiosperms, with the latter subdivided into ANA grade, *Chloranthales*, and *Magnoliids*. Taxonomic groups are labeled at the family level (lycophytes, gymnosperms and angiosperms) or order level (ferns). For each plant species, the following metrics are presented: species label, total number of detected aa-ECRTs, number of representative aa-ECRTs (aa-ECRTs rep), number of reference OTUs detected (OTU ref), number of newly identified OTUs (New OTU), genome size (in Mbp), and N50 contig value (in kbp), as computed by QUAST ^62^. One aa-ECRT, which was detected in the genome of *Anthoceros agrestis*, was confirmed as a transposable element in a subsequent analysis and is therefore indicated with an asterisk (*).

**Table 1:** Distribution of new and reference OTUs across *Caulimoviridae* clades. The table presents the distribution of newly identified and previously defined reference OTUs across the three major clades of the *Caulimoviridae* family: Clade A, Clade B, and Clade C. The classification is based on the phylogenetic reconstruction shown in Figure 4.

|  | Clade A | Clade B | Clade C |
| --- | --- | --- | --- |
| **new OTU** | 5, 6, 10, 13, 15, 21, 32 | 1, 2, 3, 4, 7, 8, 9, 11, 12, 14, 16, 17, 18, 20, 22, 23, 24, 25, 26, 27, 28, 29, 30, 31, 33, 35 | 19 |
| **ref OTU** | Badnavirus,<br>Caulimovirus,<br>Cavemovirus,<br>Dioscovirus+Badnavirus_D.alata,<br>Rosadnavirus,<br>Ruflodivirus,<br>Solendovirus,<br>Soymovirus,<br>Tungrovirus,<br>Unclassified_gp_G.max,<br>Unclassified_gp_V.vinifera+Yendovirus<br>+Orendovirus,<br>Vaccinivirus,<br>Wendovirus+Xendovirus_gp_S.tuberos<br>um,<br>Wendovirus_4,<br>Xendovirus_gp_G.raimondii,<br>Zendovirus | Fernendovirus_1_gp_C.protusa,<br>Fernendovirus_1_gp_P.formosana,<br>Fernendovirus_1_gp_P.glycyrrhiza,<br>Fernendovirus_2,<br>Gymnendovirus_1+PglaV_2,<br>Gymnendovirus_2_gp_G.biloba,<br>Gymnendovirus_2_gp_G.biloba_2+GbilV,<br>Gymnendovirus_2_gp_P.glauca+PtaeV_2+Pinus_nigra,<br>Gymnendovirus_3+PglaV_1,<br>Gymnendovirus_4_gp_P.abies,<br>Gymnendovirus_4_gp_P.glauca+PtaeV_1,<br>Petuvirus_gp_C.sinensis,<br>Petuvirus_gp_E.salsugineum,<br>Petuviruslike_C.sinensis,<br>Welwitschia_mirabilis_virus_1,<br>Welwitschia_mirabilis_virus_2 | NA |

Based on the 80% minimum amino acid (aa) identity threshold used by Caulifinder branch B ^20^, these sequences were grouped by host species into 1,684 clusters. The longest sequence from each cluster was selected as the representative aa-ECRT (aa-ECRT rep, Figure 1). The resulting set of 1,684 aa-ECRT rep was subsequently filtered to select the most divergent sequences. For this, we performed iterations of sequence clustering, alignment with 98 aa-RT reference caulimovirid sequences, and manual curation, resulting in the selection of a subset of 261 aa-ECRT rep. We also searched publicly available transcriptomic datasets (from NCBI GenBank ^23^, the 1KP project ^24^, and datasets from two transcriptomic studies dedicated to ferns and lycophytes, respectively ^25,26^) for aa-ECRTs. Using the filtering process described above yielded two additional aa-ECRTs, from the transcriptomes of the ferns *Brainea insignis* (GFUE01038349.1) and *Pteris vittata* (GGXV01034871.1), bringing the total number of aa-ECRT rep in the subset to 263.

Using an amino acid similarity threshold of 62%, allowing for clear separation of viral genera recognized by the ICTV into distinct operational taxonomic units (OTUs), the 263 aa-ECRT rep sequences combined with the 98 caulimovirid aa-RT reference sequences and eight outgroup sequences (from *Metaviridae* and *Retroviridae*) were grouped into 71 genus-level OTUs (Supplementary Figure 2; Supplementary Table 3). Thirty-six OTUs included at least one aa-RT reference caulimovirid sequence. They were named either after their corresponding ICTV-recognized genus ^6^ (e.g., OTU Soymovirus) or after previously defined OTUs ^14,16–19,27^ (e.g., OTU Florendovirus), and are referred to as “reference OTUs”.

As expected, some of the 36 reference OTUs corresponded to subdivisions of OTUs defined in previous works that used lower sequence similarity thresholds ^16,19^ (Supp. Table 3). For instance, OTU Gymnendovirus 2 ^16^ was split into three distinct OTUs, two of which included β-type gymnosperm endogenous caulimovirus-like viruses (β-GECV) ^17^. The remaining 35 OTUs lacked reference sequences and were considered novel. These mainly consisted of sequences retrieved from plant genomes or transcriptomes that had not previously been screened for aa-ECRTs. Gymnosperms proved a particularly rich source of novel OTUs, especially *Wollemia nobilis* (*Araucariaceae*), which harbored five (Figure 1).

#### Phylogenetic analysis

To determine the phylogenetic placement of the 35 novel OTUs, we performed a phylogenetic analysis using nucleotide sequences encoding the RT-RH1 domain (ntRT-RH), the most conserved region within caulimovirid genomes ^6^. Such sequences were extracted from the 55 tracheophyte genomes in the dataset that contain ECVs (Supp. Table 4) using the Caulifinder pipeline branch A, which reconstructs caulimovirid consensus sequences by aligning at least five ECV copies from the same host, sharing at least 85% identity. After filtering out LTR retrotransposons, 9,031 ECV consensus sequences were recovered from 49 plant species (three lycophytes, six ferns, and all gymnosperms (27), basal angiosperms (3), *Chloranthaceae* (1), and Magnoliids (9)) (Supp. Table 4).

Each ECV consensus sequence containing an ntRT-RH1 sequence (n=6,503) was assigned to one of the 71 previously defined OTUs, and 47 ECV consensus sequences representing 22 novel OTUs were selected. Among these, 17 OTUs were represented by two to four consensus sequences (Supp. Figure 3). For 17 of the 18 remaining novel OTUs, where either none or a single consensus sequence was available, ntRT-RH1 sequences were selected from 21 genomic ECV copies, and from one transcript from *Brainea insignis* (GFUE01038349.1) (Supp. Figure 3). In total, 69 ntRT-RH1 sequences representing 34 of the 35 novel OTUs were obtained. The only exception was OTU 34, for which only highly mutated ntRT-RH1 sequences were recovered, precluding the selection of a representative sequence (Supp. Table 5). The dataset was incremented with 73 ntRT-RH1 sequences from reference OTUs, including 40 ntRT-RH1 sequences from representative members, and 33 ntRT-RH1 sequences identified in this study from 8 angiosperm genomes (Supplementary Table 5; Supplementary Table 6). For three of the 36 reference OTUs, no representative ntRT-RH1 sequences could be recovered. Altogether, 142 ntRT-RH1 sequences representing 67 OTUs were compiled for phylogenetic analysis (Supp. Table 5).

Two phylogenetic trees were constructed using Maximum Likelihood (ML) and Bayesian methods (Supp. Figure 4), both yielding highly similar topologies. We selected the Bayesian tree for further analysis because it provided stronger support at basal nodes (posterior probability ≥ 0.95). Rooted with *Saccharomyces cerevisiae Ty3 virus* (*Metaviridae* exemplar virus), the tree was simplified by collapsing branches containing a single OTU (Figure 2). The 142 caulimovirid ntRT-RH1 sequences grouped according to their respective OTUs, except for the OTU ‘Wendovirus + Xendovirus gp S. tuberosum’ which appeared polyphyletic and the single sequence from OTU 30, which was nested within OTU 1 (Supp. Figure 4; Figure 2).

**Figure 2:**
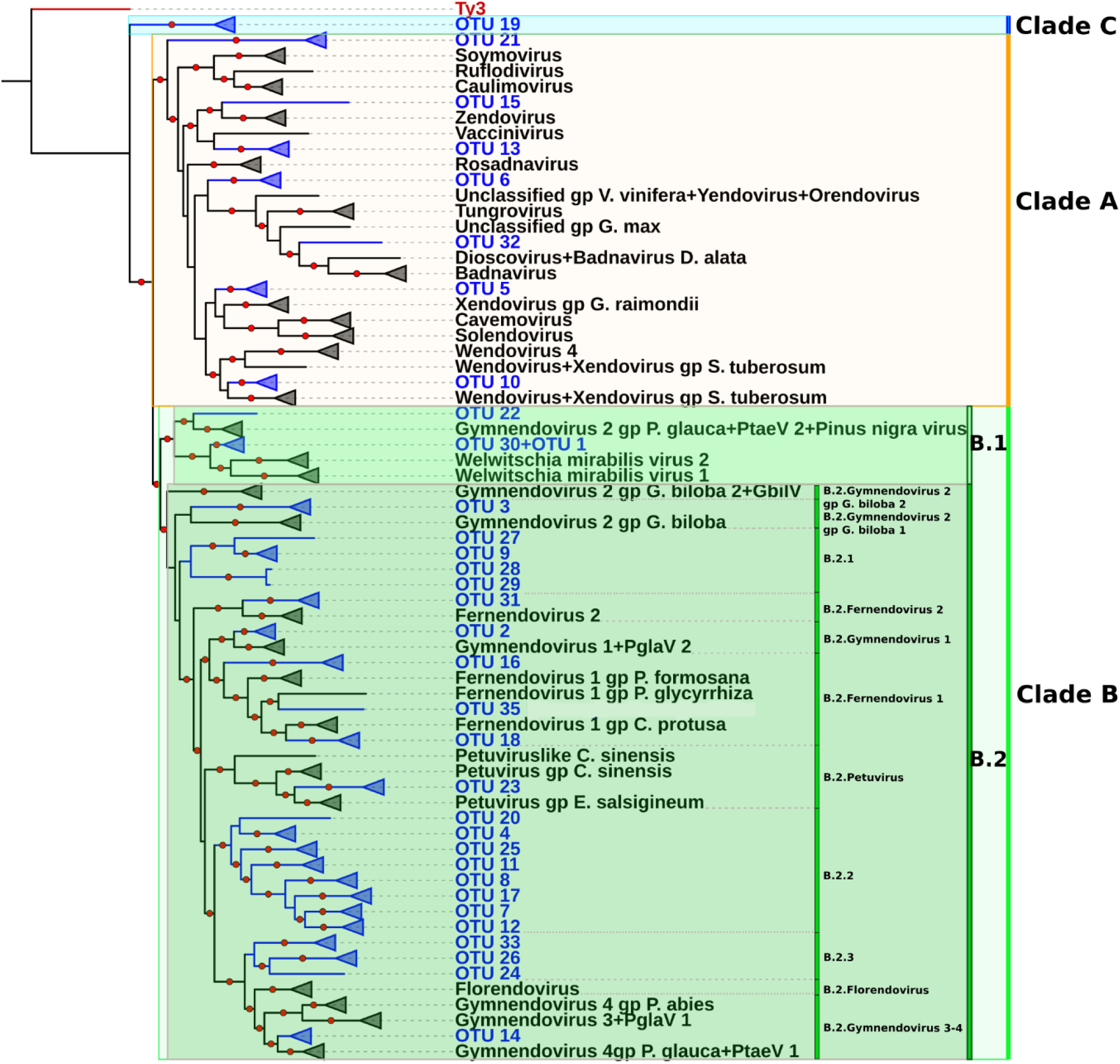
Bayesian phylogeny of *Caulimoviridae*, with collapsed OTU branches. The phylogenetic tree comprises 142 nucleotide sequences corresponding to the RT-RH1 domain of *Caulimoviridae*. Bayesian inference was conducted using MrBayes ^52–54^, and the tree is rooted with a *Metaviridae* Ty3 sequence (highlighted in red). Red circles indicate nodes with posterior probability supports ≥ 0.95. Triangles represent collapsed monophyletic groups corresponding to individual OTUs. Polyphyletic OTUs are shown in multiple locations; “Wendovirus+Xendovirus gp S. tuberosum” appears twice. OTUs newly identified in this study are highlighted in blue, while previously defined (reference) OTUs are shown in black. The tree is divided into three major clades, color-coded as follows: Clade A (orange), Clade B (green), and Clade C (blue).

**Figure 3:**
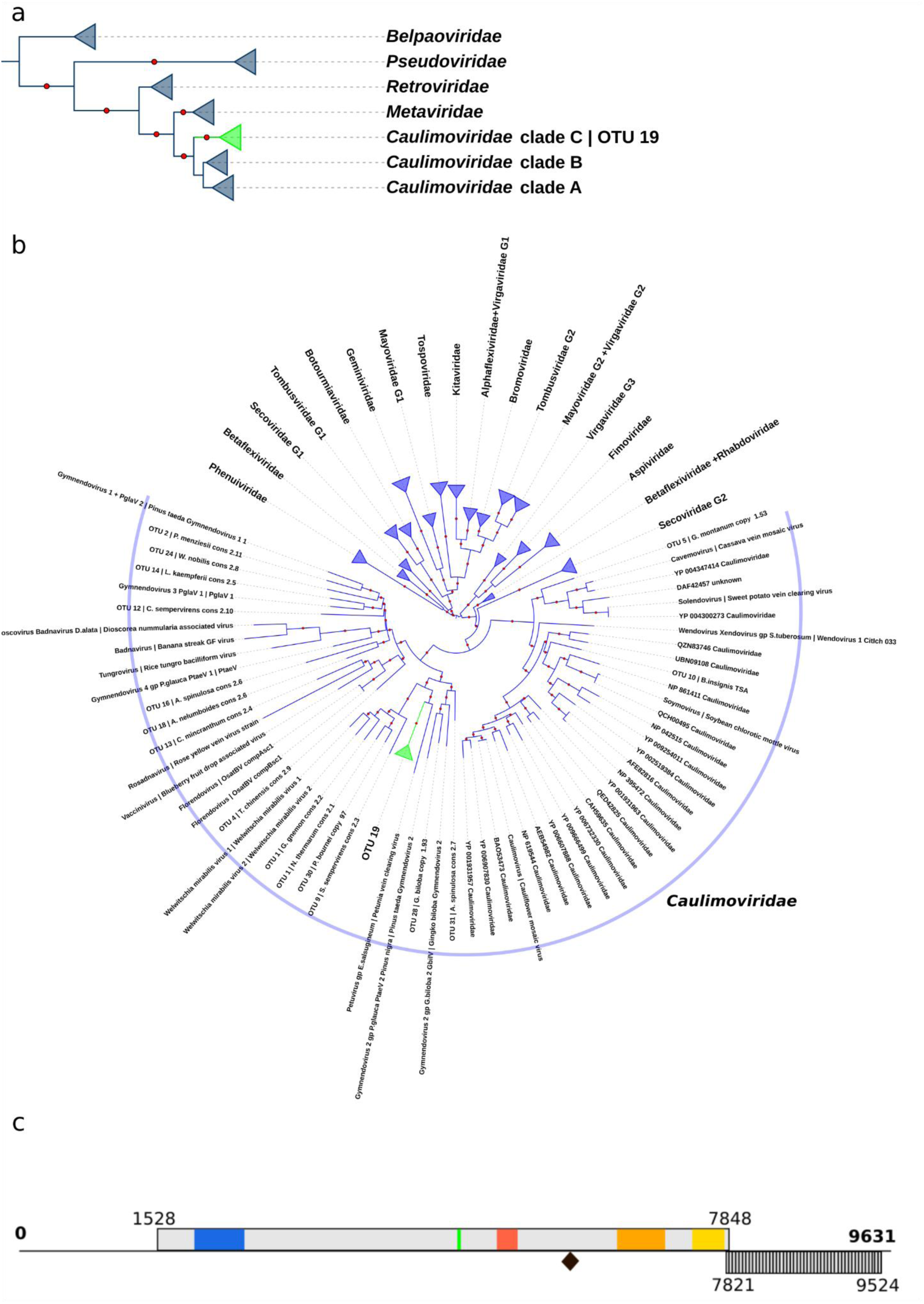
Taxonomic placement of Clade C and genome structure of Wollendoviruses. (a) ML phylogeny of reverse transcriptase domains from 52 *Ortervirales* sequences. Red dots mark branches with bootstrap support ≥ 50%. Clades are collapsed for *Caulimoviridae* and by family for other *Ortervirales* members. (b) ML phylogeny of the 30K movement protein (MP) family, including 46 *Caulimoviridae* sequences identified using Caulifinder and 286 sequences from Butkovic *et al.* (2024) ^29^, representing the following plant viral families: *Alphaflexiviridae*, *Aspiviridae*, *Betaflexiviridae*, *Bromoviridae*, *Botourmiaviridae*, *Caulimoviridae*, *Fimoviridae*, *Geminiviridae*, *Kitaviridae*, *Mayoviridae*, *Phenuiviridae*, *Rhabdoviridae*, *Secoviridae*, *Tospoviridae,* and *Virgaviridae*. The tree is midpoint-rooted. For *Caulimoviridae* sequences, OTU IDs are listed, followed by sequence names; NCBI accession numbers are provided for reference sequences. OTU 19 is highlighted in green. Red dots indicate branches supported by bootstrap values >= 80 %. For non-caulimovirid families, branches are collapsed by family and represented as triangles. (c) Genome organisation of the proposed Wollendovirus type member, WolV1, reconstructed from sequence assembly. Open reading frames (ORFs), predicted using ORF Finder (https://www.ncbi.nlm.nih.gov/orffinder/), are shown as light grey rectangles. ORF of unknown function is shaded. The black diamond indicates the tRNAMet primer binding site, corresponding to genome position +1. Conserved domains identified via CD-search (https://www.ncbi.nlm.nih.gov/Structure/cdd/wrpsb.cgi) against the CDD database ^57^ are color-coded: viral movement protein (MP; PF01107) in blue; retropepsin (AP; CD00303) in red; reverse transcriptase (RT; CD01647) in orange; RNase H1 (RH1; CD06222) in yellow; zinc finger in green.

**Figure 4:**
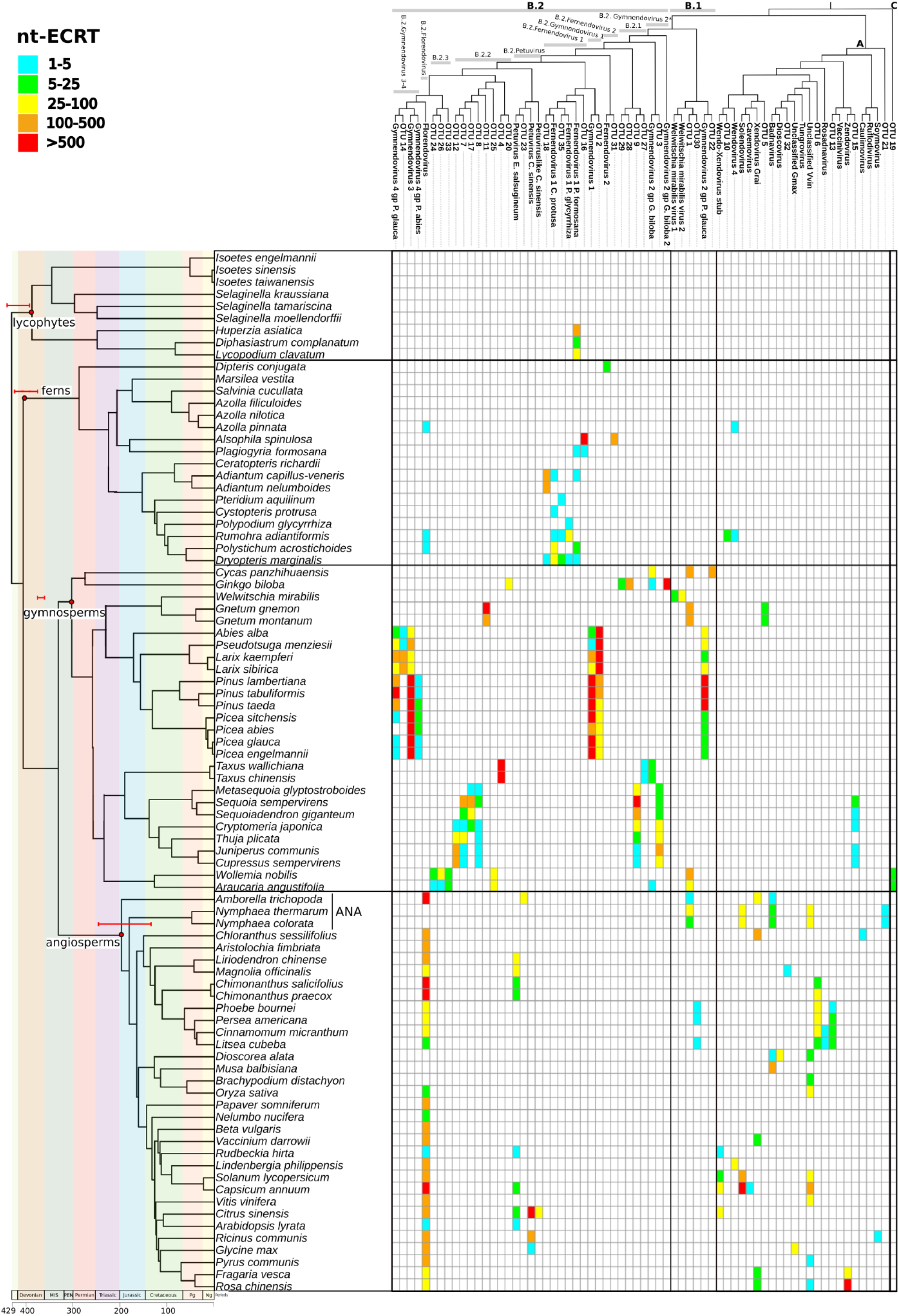
**Heatmap of nt-ECRT abundance by OTU across plant hosts** The heatmap displays the distribution of nt-ECRTs across plant genomes, grouped by host species (left) and viral OTUs (top right). Left: the host plant phylogeny is adapted from the TimeTree ^63^ database (accessed on 20/09/24). Geological periods are shown along the top, with red bars indicating the standard deviation of estimated emergence times for major plant lineages, based on the following sources: Lycophytes (432.5–392.8 MYA; Morris *et al.*, 2018 ^38^), Ferns (411.5- 384 MYA Morris *et al.*, 2018 ^38^; Lehtonen *et al.*, 2017 ^37^), Gymnosperms (380–360 MYA Rothwell *et al.*, 1989 ^64^; Stull *et al.*, 2021 ^28^), and Angiosperms (246.5–130 MYA; Herendeen *et al.*, 2017 ^35^). Top: the *Caulimoviridae* cladogram is derived from the Bayesian tree of 142 RT-RH1 nucleotide sequences (see Figure 4), with major clades color-coded as follows: Clade A (orange), Clade B (green), and Clade C (blue). For visualization purposes, the polyphyletic OTU ‘Wendovirus+Xendovirus gp S. tuberosum’ (Figure 4) is represented here as a single, monophyletic clade. For clarity some OTU names have been shortened: Xendovirus gp G. raimondii to Xendovirus Grai; Unclassified gp V. vinifera+Yendovirus+Orendovirus to Unclassified Vvin; Dioscovirus+Badnavirus D. alata to Dioscovirus; Wendovirus+Xendovirus gp S. tuberosum to Wendo-Xendovirus stub; Gymnendovirus 2 gp P. glauca+PtaeV 2+Pinus nigra virus to Gymnendovirus 2 gp P. glauca; Gymnendovirus 2 gp G. biloba 2 + GbilV to Gymnendovirus 2 gp G. biloba 2; Gymnendovirus 1 + PglaV2 to Gymnendovirus1. Color intensity in the heatmap corresponds to the number of nt-ECRT copies assigned to each OTU in a given host genome.

The tree is divided into three main clades, A, B, and C, with Clades A and B previously defined by Diop *et al.* (2018) ^16^ and Clade B subdivided into two subclades: B.1 and B.2. Seven of the 34 new OTUs (Table 1) grouped within Clade A. Among these, OTU 21 was sister to all Clade A members, while the other six were closer to either previously described ECVs or ICTV-recognized genera. In Clade B, 26 novel OTUs (Table 1) were distributed across the topology; half of them were closely related to known ECVs while twelve formed two new groups, B.2.1 and B.2.2. In addition, OTU 23, was grouped with *Petuvirus*, the only ICTV-recognized genus in Clade B. Subclade B.2 was further divided into groups of closely related OTUs to better describe its extended diversity (Figure 2). Remarkably, our phylogenetic analysis placed OTU 19 as a sister to Clades A and B, leading to its provisional assignment to a new Clade C.

#### Characterization of caulimovirid Clade C

In both unrooted ML and Bayesian trees (Supp. Figure 4), OTU 19 formed a polytomy with the outgroup (Ty3) and *Caulimoviridae* Clades A and B, leaving its placement unresolved. To clarify its phylogenetic position within *Ortervirales*, we performed an ML phylogenetic analysis using RT domain amino acid sequences (aa-RT) from OTU 19, alongside representatives of *Caulimoviridae* Clades A and B and of the four other *Ortervirales* families (Supp. Table 7), following the RT-based framework of Krupovic *et al.* (2018) ^7^.

The resulting tree resolved the polytomy (Figure 3a), placing OTU 19 as a sister clade to Clades A and B. OTU 19 includes sequences from *Wollemia nobilis* and *Araucaria angustifolia*, two critically endangered *Araucariaceae* species ^28^. The 30K MP domain is shared among many viruses, prompting us to analyze the phylogenetic relationships of the MP domains from OTU 19 sequences alongside those from other caulimovirid clades and 14 plant virus families ^29^. While MPs from many families were polyphyletic, *Caulimoviridae* MPs formed a monophyletic group with OTU 19 nested within (Figure 3b).

We assembled a representative genome from *W. nobilis* ECVs, and the corresponding virus was tentatively named Wollendovirus 1 (WolV1; Figure 3c). The WolV1 genome spans 9,631 bp and contains a 16 bp tRNAMet (TGGTATCAGAGCCAGG) primer binding site, unique in its genomic position but analogous to other caulimovirids ^6^. WolV1 contains two putative ORFs encoding respectively a large polyprotein (2,107 aa, 243 kDa) featuring canonical caulimovirid domains (MP, zinc finger (ZnF) typical of caulimovirid capsid proteins, AP, RT and RH1), and a 568 aa (67 kDa) protein of unknown function with no homolog in GenBank.

The substantial structural and organisational similarity between the WolV1 genome and those of known caulimovirids suggests a synapomorphic relationship. Combined with the close phylogenetic relationship between OTU 19 MP and RT domains with those of other caulimovirids, this evidence supports the classification of Clade C as a distinct new lineage within the *Caulimoviridae* family.

#### Distribution of ECVs across plant genome assemblies

To assess the distribution of ECVs across plant taxa, we screened 93 land plant genome assemblies (73 previously analyzed plus 20 monocot and dicot assemblies to extend our analysis to most of the angiosperm groups; Supp. Tables 6, 8) for nucleotide sequences of endogenous caulimovirid RTs (nt-ECRTs). Using nucleotide sequences enabled the inclusion of sequences with mutations that disrupt reading frames. From this analysis, we identified 47,135 nt-ECRTs across 75 tracheophyte genomes (Supp. Table 9). Copy numbers were systematically higher than for aa-ECRTs, averaging threefold and reaching up to 18.7-fold in *Metasequoia glyptostroboides*. No nt-ECRTs were detected in genomes lacking aa-ECRTs. The abundance of nt-ECRTs was positively correlated with the host genome size (r^2^ = 0.45, p=9e^-13^). This trend held within lycophytes (r^2^ = 0.85, p=4e^-04^), gymnosperms (r^2^ = 0.46, p=2.8e^-04^), and angiosperms (r^2^ = 0.43, p=0.008) after multiple testing correction ^30^, but was weaker in ferns (r^2^ = 0.24, p=0.03) (Supp. Figure 5).

**Figure 5:**
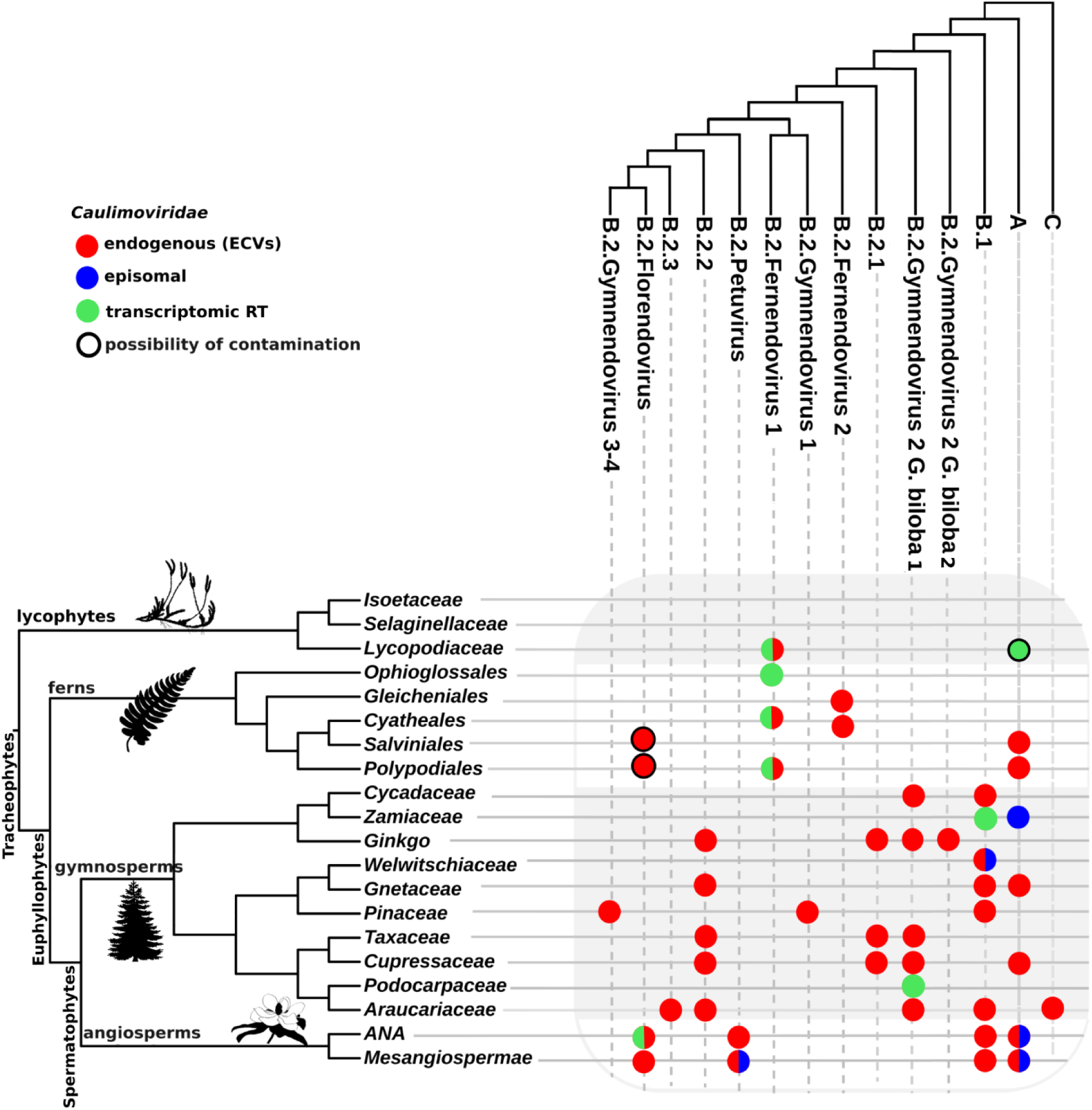
**Distribution of *Caulimoviridae* clades and subclades across plant taxa** Left: a plant cladogram depicts the major plant divisions of embryophytes for which either genomic or transcriptomic data were analyzed. Lycophytes and gymnosperms are resolved to the family level due to broader data availability. Ferns, with more limited genomic representation, are shown at the order level. Angiosperms are split into basal ANA grade and *Mesangiospermae*, which includes *Magnoliids*, *Chloranthales*, monocots, and dicots. Top: A viral cladogram summarizes the major *Caulimoviridae* clades and subclades as defined in Figure 3. Intersection circles indicate the detection of *Caulimoviridae* sequences within each host taxon, with the following color code. Blue: detection of episomal viral forms, including Petunia vein clearing viruses (B.2 Petuvirus ^6^), Welwitschia mirabilis viruses 1 and 2 ^18^, cycad leaf necrosis virus (Clade A, genus *Badnavirus* ^40^), and additional ICTV-recognized *Caulimoviridae* ^6^. Red: detection of ECVs in host genomes. Green: detection of caulimovirid transcripts in transcriptomic datasets. Circles combining multiple colors indicate multiple forms of evidence (E.g., genomic + transcriptomic). Silhouettes are sourced from the PhyloPic image repository (https://www.phylopic.org/).

All nt-ECRTs were assigned to OTUs using BLASTx against representative aa-RTs with a stringent 65% amino acid identity threshold, which we chose slightly above the OTU demarcation threshold (62%) to ensure unambiguous assignment (Figure 4, Supp. Table 10). All 47,135 nt-ECRTs were assigned to an OTU: 32,303 were assigned to 33 reference OTUs and 14,832 to 34 novel OTUs. Distribution varied widely between OTUs in both sequence abundance and host plant diversity. For example, *Capsicum annuum* contained the highest number of nt-ECRTs linked to a single OTU, with 1,945 nt-ECRTs assigned to *Solendovirus* (Figure 4, Supp. Table 10). Florendoviruses exhibited the broadest host range, with 6,982 nt-ECRTs identified across 28 angiosperm species. In contrast, OTU 21 was represented by only two nt-ECRTs, in ANA-grade species *Nymphaea colorata* and *Nymphaea thermarum*.

Strong lineage-specific patterns were observed across the three caulimovirid clades. In Clade A, nt-ECRTs were predominantly detected in angiosperm genomes (Figure 4) with four notable exceptions: (i) OTU 5 was restricted to *Gnetaceae* (*Gnetum gnemon* and *Gnetum montanum*); (ii) OTU 15 was detected solely in five of seven *Cupressaceae* genomes (*Sequoia sempervirens*, *Sequoiadendron giganteum*, *Cryptomeria japonica*, *Juniperus communi*s and *Cupressus sempervirens*); (iii) Wendovirus 4-related nt-ECRTs were found in both angiosperms (*Lindenbergia philippensis*) and ferns (*Azolla pinnata* and *Rumohra adiantiformis*); and (iv) OTU 10 was restricted to the fern *Rumohra adiantiformis*, with transcriptomic evidence suggesting its presence in the ferns *Brainea insignis* and *Pteris vittata*. Furthermore, OTU 21, which occupied a basal position within Clade A, was limited to the basal angiosperm family *Nymphaeaceae*. In Clade B, nt-ECRTs were primarily restricted to lycophyte, fern, and gymnosperm genomes, with two notable exceptions: Florendovirus and Petuvirus were exclusive to angiosperms and frequently found in high copy numbers. Additionally, OTU 1 (subclade B.1) exhibited a broad host range, encompassing gymnosperms and basal angiosperms (*Nymphaeaceae, Amborellaceae)*. In contrast, its sister lineage, OTU 30, was restricted to magnoliids. Finally, Clade C (OTU 19) nt-ECRTs were strictly host-specific, detected only in *Araucariaceae*, specifically in *Wollemia nobilis* and *Araucaria angustifolia*. We also detected OTU 19 sequences in genomic reads from a third *Araucariaceae* species, *Agathis dammara*.

#### Identification of caulimovirid sequences in plant transcriptomes

Transcriptomic analysis can reveal the presence of OTUs in host species, whether transcripts derive from ECVs or from episomal caulimovirids. To further investigate caulimovirid-host interactions in basal plant lineages, we used Caulifinder branch B ^20^ on the same transcriptomic dataset as above ^23–26^.

We identified 529 transcripts encoding caulimovirid RT domains, 491 of which originated from angiosperms (Supp. Table 11). For taxonomic assignment, we selected 194 transcripts covering ≥ 80% of the RT domain. Notably, 13 transcripts were from lineages absent in genome datasets: eight from ferns, two from gymnosperms, and three from ANA-grade angiosperms (Table 2; Supp. Figure 6). Fern sequences originated from the orders *Cyatheales*, *Polypodiales*, *Marattiales*, and *Ophioglossales*, the latter two representing basal lineages ^31^. One of these sequences was assigned to the OTU Fernendovirus 1 P. formosana and two to the OTU Fernendovirus 1 C. protusa, within the ‘B.2.Fernendovirus 1’ group (Figure 2, Table 2). The remaining five fern sequences lacked sufficient identity (below 62%) for OTU assignment using tBLASTn, but two, from *Botrypus virginianus* (*Ophioglossales*) and *Lindsaea linearis* (*Polypodiales*), placed within the ‘B.2.Fernendovirus 1’ group (Supp. Figure 7). The two gymnosperm transcripts, from families *Zamiaceae* (*Cycadales*) and

**Figure 6:**
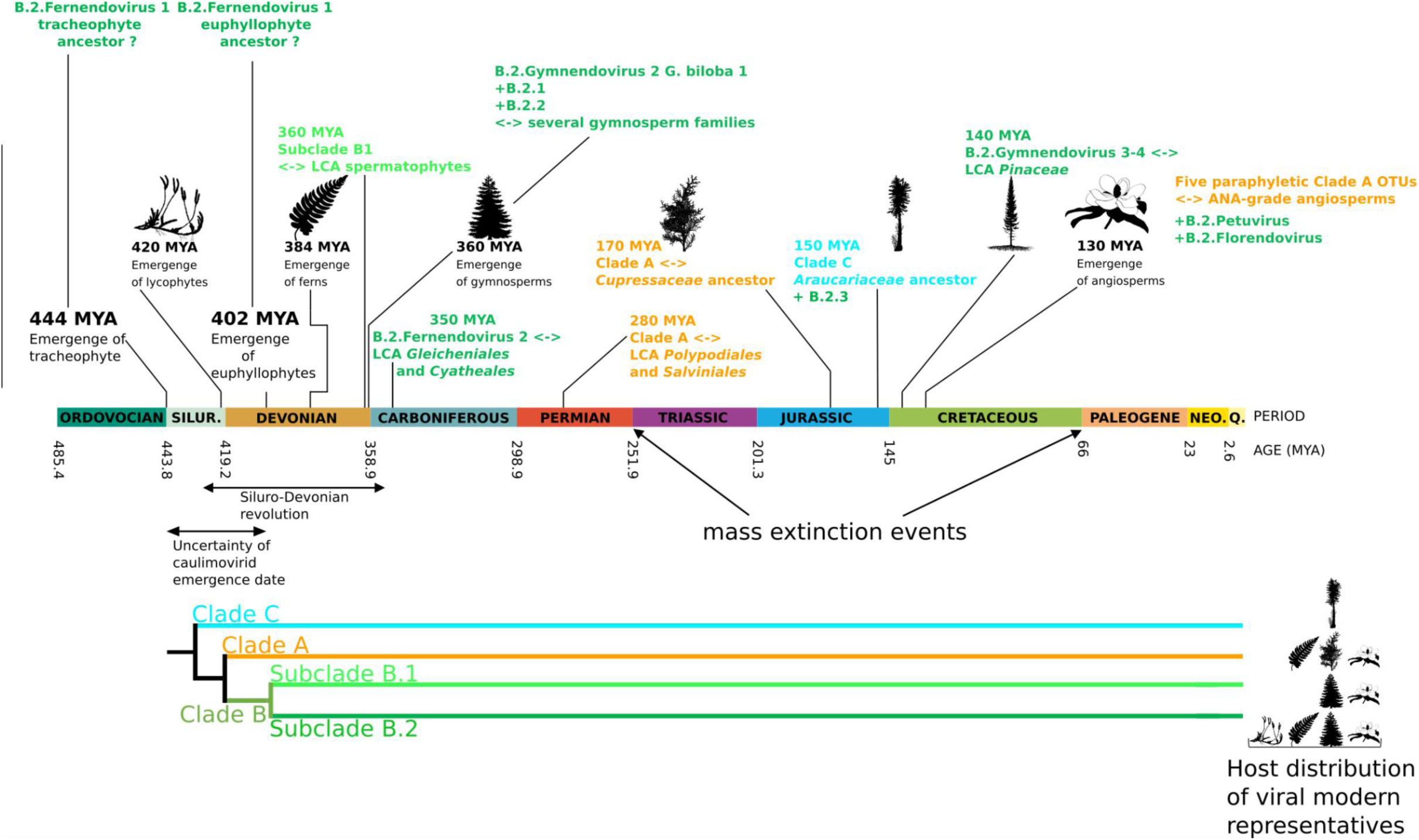
**Evolution of *Caulimoviridae* diversity over time in relation to host range** The figure integrates viral diversity with plant evolutionary history along a geological timescale adapted from McLoughlin, (2021) ^39^. Timeline: key evolutionary milestones are indicated, including (i) the emergence of tracheophytes and euphyllophytes ^38^; (ii) the divergence of major vascular plant lineages: lycophytes, ferns, gymnosperms and angiosperms ^28,35–38^; (iii): the last common ancestor (LCA) of *Gleicheniales*-*Cyatheales* and *Polypodiales*-*Salviniales* ^37^; and (iv) the LCA *Cupressaceae* and *Araucariaceae* ^28^. The known host range of each clade is summarized and matched to key plant divergence points on the geological timeline, illustrating the potential cospeciation and diversification of *Caulimoviridae* alongside their plant hosts. Bottom: A simplified *Caulimoviridae* cladogram (adapted from Figure 4) highlights the three main viral clades: A (orange), B (green), and C (blue). Silhouettes representing plant taxa are sourced from the PhyloPic image repository (https://www.phylopic.org/).

**Table 2:** nt-RTs detected in the three transcriptome sources. This table presents details of the 18 transcripts retained from the search for caulimovirid RTs in plant transcriptomes. Each entry includes the following information : (i) Taxonomic information of the plant: division (column 1), species (column 2), genus (column 3), family (column 4); (ii) BLAST analysis results against the 361 aa-RT sequences used to define OTUs: OTU with the best hit (column 5), identity percentage of the best hit (column 6), transcript length in bp (column 7); (iii) Reference information: source publication (column 8), transcript accession number (column 9).

| Plant division | Species | Genus | Family | OTU best hit | identity | size (nt) | Origin | Accession |
| --- | --- | --- | --- | --- | --- | --- | --- | --- |
| Lycophytes | *Lycopodium annotinum* | *Lycopodiales* | *Lycopodiaceae* | Fernendovirus 1<br>*P. formosana* | 76.2 % | 2028 | 1kp | scaffold-ENQF-2084799-Lycopodium |
|  | *Pseudolycopodiella caroliniana* | *Lycopodiales* | *Lycopodiaceae* | Badnavirus | 72.6 % | 540 | 1kp | scaffold-UPMJ-2015667-Pseudolycopodiella |
|  | *Palhinhaea hainanensis* | *Lycopodiales* | *Lycopodiaceae* | Badnavirus | 67.2 % | 1362 | https://doi.org/10.1016/j.plid.2021.08.004 | Palha_Gene.73649_Cluster-24281.0 |
|  | *Palhinhaea hainanensis* | *Lycopodiales* | *Lycopodiaceae* | 35 | 62.9 % | 804 | https://doi.org/10.1016/j.plid.2021.08.004 | Palha_Gene.103642_Cluster-38429.0 |
|  | *Lycopodium zonatum* | *Lycopodiales* | *Lycopodiaceae* | Fernendovirus 1<br>*P. formosana* | 65.7 % | 1128 | https://doi.org/10.1016/j.plid.2021.08.004 | Lyzo_Gene.43739_c45295_g1_i1 |
| Ferns | *Botrypus virginianus* | *Ophioglossales* | *Ophioglossaceae* | Fernendovirus 1<br>*P. formosana* | 47.8 % | 2791 | 1kp | scaffold-BEGM-2004510-Botrypus |
|  | Unclassified Marattia | *Marattiales* | *Marattiaceae* | 31 | 56.1 % | 166 | 1kp | scaffold-UXCS-2117714-Marattia |
|  | *Sceptridium dissectum* | *Ophioglossales* | *Ophioglossaceae* | 25 | 48.2 % | 3425 | 1kp | scaffold-EEAQ-2089744-Sceptridium |
|  | *Lindsaea linearis* | *Polypodiales* | *Lindsaeaceae* | Fernendovirus 1<br>*P. formosana* | 50.2 % | 2371 | 1kp | scaffold-NOKI-2097008-Lindsaea |
|  | *Lonchitis hirsuta* | *Polypodiales* | *Lonchitidaceae* | Fernendovirus 1<br>*C. protusa* | 79 % | 6523 | 1kp | scaffold-VVRN-2010621-Lonchitis |
|  | *Amblovenatum opulentum* | *Polypodiales* | *Thelypteridaceae* | Fernendovirus 1<br>P. glycyrrhiza | 55.2 % | 465 | https://doi.org/10.1101/2024.08.27.609851 | Aop_g40581 |
|  | *Cibotium barometz* | *Cyatheaales* | *Cibotiaceae* | Fernendovirus 1<br>C. protusa | 77.9 % | 483 | https://doi.org/10.1101/2024.08.27.609851 | Cba_g18761 |
|  | *Nephrolepis biserrata* | *Polypodiales* | *Nephrolepidaceae* | Fernendovirus 1<br>P. formosana | 67.9 % | 1362 | https://doi.org/10.1101/2024.08.27.609851 | Nbi_g42169 |
| **Gymnosperms** | *Dioon edule* | *Cycadales* | *Zamiaceae* | 1 | 71 % | 509 | 1kp | scaffold-WLIC-2000696-Dioon |
|  | *Podocarpus rubens* | *Pinales* | *Podocarpaceae* | 3 | 62.3 % | 6552 | 1kp | scaffold-XLGK-2060667-Podocarpus |
| **ANA** | *Austrobaileya scandens* | *Austrobaileyaales* | *Austrobaileyaceae* | Florendovirus | 79.3 % | 1747 | 1kp | scaffold-FZJL-2010292-Austrobaileya |
|  | *Austrobaileya scandens* | *Austrobaileyaales* | *Austrobaileyaceae* | Florendovirus | 66 % | 4766 | 1kp | scaffold-FZJL-2022942-Austrobaileya |
|  | *Austrobaileya scandens* | *Austrobaileyaales* | *Austrobaileyaceae* | Florendovirus | 78.5 % | 583 | 1kp | scaffold-FZJL-2162197-Austrobaileya |

*Podocarpaceae* (*Araucariales*), were assigned to OTUs 1 and 3, respectively. For ANA-grade angiosperms, three transcripts from *Austrobaileya scandens* (*Austrobaileyales*) were classified as Florendovirus, two clustering near OsatBV in the ‘B.2 Florendovirus’ group (Supp. Figure 8) ^14^. Additionally, five transcripts from *Lycopodiaceae*, an underrepresented family in genome datasets, were analyzed. Two were assigned to OTU Fernedovirus 1 P. formosana and one to OTU 35 within the ‘B.2.Fernendovirus 1’ group, while two clustered with Badnavirus sequences from angiosperms.

#### Evidence for patterns of cospeciation

To investigate the macroevolution of *Caulimoviridae*, we integrated host-virus associations from genomic and transcriptomic data, as well as from episomal caulimovirids. Figure 5 illustrates a condensed interactome linking viral clades or subclades to plant families or orders. In combination with the distribution data (Figure 4), this synthesis enabled us to investigate the patterns of caulimovirid-host coevolution at a macroscopic scale. We found that caulimovirids and their hosts do not exhibit overall parallel phylogenies, some plant divisions harboring distinct caulimovirid lineages, such as Clade A, B and C being associated with gymnosperm hosts (Figures 4 & 5). However, we observed several host-caulimovirid distribution subpatterns that align with the coexistence of multiple deeply rooted caulimovirid lineages (Figure 6), outlined in the following points i) through v).

i. **Clade C** sequences, all grouped within OTU 19, were found exclusively in three extant *Araucariaceae* species - *W. nobilis*, *A. angustifolia*, and *A. dammara* (the latter being based on SRA data) - each representing one of the family’s three living genera (Figure 4). These species are distributed across Australia, South America, and South East Asia, regions that became isolated following the breakup of Gondwana, which concluded ∼100 MYA during the Cretaceous ^32,33^. Molecular dating places their last common ancestor (LCA) at least 150 MYA28.

The narrow host range of Clade C, aligned with *Araucariaceae* biogeography, suggests their LCA hosted episomal Clade C viruses that co-diverged within the family. This supports the idea that Clade C ECVs represent descendants of a *Caulimoviridae* lineage that originated no later than the mid-Jurassic and evolved alongside their slow-evolving hosts^34^. The aa-ECRTs of Clade C show high levels of pairwise identity, up to 81.3%, 82.9% and 84% between *W. nobilis* and *A. angustifolia*, *A. angustifolia* and *A. dammara*, and *W. nobilis* and *A. dammara*, respectively. Such conservation over long timescales may reflect the slow evolution of both the virus and its host, potentially due to strong selective constraints.

(ii) **Clade A** sequences were predominantly found in angiosperms. Yet, six paraphyletic OTUs (*Solendovirus*, Xendovirus gp G. raimondii, *Badnavirus*, *Caulimovirus*, Unclassified gp V. vinifera, and OTU 21) also occurred in ANA-grade lineages (Figure 4). In addition, OTU 21, restricted to *Nymphaeaceae*, occupies a basal position in Clade A, suggesting that diversification of this clade predated or coincided with early angiosperm evolution, ∼ 130 MYA ^35,36^.

The host range of some Clade A OTUs extends beyond angiosperms. For example, OTU 15 spanned five *Cupressaceae* species, including *Cupressus sempervirens* and *Sequoia sempervirens*, whose LCA dates to at least 170 MYA ^28^. Two closely related OTUs (OTU 10 and Wendovirus 4) were detected in ferns of the orders *Polypodiales* (*Brainea insignis*, *Pteris vittata*, *Rumohra adiantiformis*) and *Salviniales* (*Azolla pinnata*), which diverged over 280 MYA, predating angiosperms ^37^. In addition, OTU 5 ECVs were found in two *Gnetaceae* species (Figure 4), and Badnavirus transcripts were found in *Lycopodiaceae* (Figure 5).

(iii) **Subclade B.1** sequences were identified across a wide phylogenetic range of seed plants (spermatophytes) hosts (Figure 5), encompassing several Magnoliid species, two ANA-grade basal angiosperm clades, and multiple distantly related gymnosperm families: *Araucariaceae*, *Gnetaceae*, *Pinaceae*, *Welwitschiaceae*, and the basal lineages *Cycadaceae* (*Cycas panzhihuaensis*) and *Zamiaceae* (*Dioon edule,* based on transcriptomic evidence, Table 2). These taxa trace back to an LCA of spermatophytes estimated at ∼360 MYA ^28^. The widespread distribution of subclade B.1 across both early-diverging gymnosperms and basal angiosperms suggests long-term cospeciation with spermatophytes, without significant transmission into more recent angiosperms.

Remarkably, sequences from Welwitschia mirabilis virus 1 and 2 (WMV1 and WMV2), derived from RNA-seq data ^18^, clustered with subclade B.1. This clustering aligns with the work of Debat & Bejerman ^18^, who suggested the existence of episomal forms of both viruses, providing indirect evidence for ongoing subclade B.1 infections in gymnosperms. This suggests that subclade B.1 includes both endogenous and episomal forms that have persisted since the early evolution of spermatophytes.

(iv) **Subclade B.2** comprises three basally branching, paraphyletic OTU groups associated exclusively with gymnosperms, potentially representing ancient *Caulimoviridae* lineages (Figure 4): (i) the ‘B.2.Gymnendovirus 2 gp G. biloba 2’ group, found only in *Ginkgo biloba*; (ii) the ‘B.2.Gymnendovirus 2 gp G. biloba 1’ group, with a broader host range spanning *Araucariaceae*, *Cycadaceae*, *Ginkgoaceae*, *Cupressaceae*, *Taxaceae* and *Podocarpaceae* (*Podocarpus rubens,* from transcriptomic data, Table 2); (iii) the B.2.1 group comprising four novel OTUs (9, 27, 28 and 29) identified in *Ginkgo biloba* and several *Cupressaceae* species. The distribution of these three B.2 groups across deeply divergent gymnosperm lineages, including “living fossils” like *Ginkgo biloba*, suggests they may be remnants of viral clades that arose early in gymnosperm evolution, possibly at the origin of seed plants ∼360 MYA. Their persistence in distinct families may reflect lineage-specific retention and extinction of related viruses, or highly constrained co-evolution with slow-evolving host lineages.
(v) In contrast to the groups in point (iv), the remaining members of subclade B.2 show a broader distribution across tracheophyte lineages (Figure 4), suggesting an older origin and complex diversification. The ‘B.2.Fernendovirus 2’ group is the most basal in subclade B.2, found in distantly related ferns *Alsophila spinulosa* (*Cyatheales*) and *Dipteris conjugata* (*Gleicheniales*), whose LCA dates back at least 350 MYA ^37^. Since ‘B.2.Fernendovirus 2’ appears absent outside ferns, there is little support for a recent cross-division host switch, though host switches between ferns remain plausible. This basal group is sister to the rest of subclade B.2, which divides into two monophyletic groups.

The first monophyletic group includes ‘B.2.Gymnendovirus 1’, found in *Cupressaceae*, and ‘B.2.Fernendovirus 1’, present in various fern orders, including the basal *Ophioglossales*, and in lycophytes. The presence of ‘B.2.Fernendovirus 1’ in ferns and lycophytes — two ancient vascular plant lineages — suggests it may have originated near the base of vascular plants (∼444 MYA ^38^).

The second monophyletic group has two branches. One includes ‘B.2.Petuvirus’, found in ANA-grade angiosperms, magnoliids, and eudicots, and ‘B.2.2’, which spans divergent gymnosperm families, including *Ginkgoaceae*, *Gnetaceae*, *Cupressaceae*, and *Araucariaceae*, whose LCA dates back at least 350 MYA ^28,39^. This broad host range suggests multiple ancient host jumps or a deep caulimovirid lineage that diversified alongside seed plants. The other branch includes ‘B.2.3’, found only in *Araucariaceae*, indicating a narrow and potentially ancient family-specific association, ‘B.2.Florendovirus’, widespread in basal angiosperms and mesangiosperm, consistent with extensive diversification within flowering plants, and ‘B.2.Gymnendovirus 3-4’, restricted to *Pinaceae*, suggesting a conifer-specific lineage.

Overall, the distribution of B.2 caulimovirids across ferns, lycophytes, gymnosperms, and angiosperms suggests a mosaic of ancient cospeciation, lineage-specific persistence, and probable host-switching, highlighting the deep and complex macroevolutionary history of this *Caulimoviridae* subclade.

## Discussion

Our study substantially advances understanding of *Caulimoviridae* diversity by identifying 35 new genus-level OTUs. Of these, 34 enrich the known diversity within Clades A and B, and one defines a previously unrecognized lineage, Clade C. For context, the ICTV currently recognizes 11 genera in the *Caulimoviridae* family, based on episomal genome structure and phylogeny ^6^. By greatly expanding the known genetic diversity of the family, our findings position the *Caulimoviridae* as a powerful model for exploring the long-term evolutionary dynamics of plant viruses.

We also show that *Caulimoviridae* have colonized all major divisions of tracheophytes, greatly broadening their recognized host range. Prior to this work, episomal and endogenous caulimovirids had been reported in 158 plant species from 68 distinct families and 59 orders ^6,14–19,21,27^ (Supp. Table 12; Supp. Table 13). Here, we detect ECVs and/or ECV transcripts in 302 plant species, expanding the total known host range to 421 plant species across 135 families and 78 orders (Supp. Table 14).

Within this expanded diversity, Clade A OTUs are found primarily in angiosperms, but also in gymnosperms and ferns. Clade B OTUs span the entire tracheophyte spectrum, uniquely including lycophytes. Whereas only a single caulimovirid reverse transcriptase sequence had been previously identified from lycophytes, our detection of ECRTs across multiple lycophyte genomes and transcriptomes confirms that lycophytes are, or have been, genuine hosts of the *Caulimoviridae*. In contrast, Clade C appears narrowly restricted to the *Araucariaceae*, suggesting a lineage-specific association. Importantly, plant genome resources remain unevenly distributed across tracheophytes, particularly in lycophytes, ferns, gymnosperms, and basal angiosperms, which likely obscures additional lineage breadth.

To investigate macroevolutionary patterns, we initially posited (a) that *Caulimoviridae* evolution was shaped by cospeciation with plant hosts, and (b) that caulimovirid have relatively narrow host ranges, typically restricted to one or a few closely related families. The latter assumption is consistent with ICTV-recognized genera, which are typically limited to one or a few related plant families rather than spanning broadly across multiple families, except for *Badnavirus*, whose broad host range likely reflects historical host switching ^6,40^. The distribution of ECVs from each OTU across plants (Figure 4) broadly supports a predominantly narrow host range, with notable exceptions such as Florendovirus, which spans all angiosperms.

Under (a) and (b), plant diversification would drive viral diversification, leading to largely parallel host and virus phylogenies, as also demonstrated at the scale of an island ecological community by French *et al.* (2023) ^41^, who showed that host phylogeny strongly shapes viral transmission networks and contributes to congruent host-virus evolutionary patterns. However, the distribution of *Caulimoviridae* lineages does not support strict cospeciation from a single ancestral lineage (Figures 4 and 5). Both Clades A and B occur in ferns, gymnosperms, and angiosperms, and several “orphan” lineages are found in basal gymnosperms. Instead, we identified several virus-host distribution subpatterns consistent with the ancient origin of multiple caulimovirid lineages (Figure 6). We interpret association of closely related OTUs with closely related hosts (at the plant division level) as evidence of vertical transmission extending back to the host LCA. In contrast, associations spanning distantly related hosts likely reflect host switches or incomplete lineage sorting (ILS).

Within this framework, Clade A cospeciation can be traced back at least to early angiosperm evolution ∼130 MYA ^35,36^. Its sporadic presence in ferns and gymnosperms may represent a deeper origin with differential retention, multiple host switches, or ILS. Subclade B.1 can be traced back to the LCA of seed plants (at least 360 MYA ^28^). Subclade B.2 is especially notable for its broadest host range, spanning angiosperms, gymnosperms, ferns, and lycophytes. The ‘B.2 Fernendovirus 1’ lineage, which includes ECVs from both ferns and lycophytes (Figure 6), is particularly informative for reconstructing early virus-plant associations. Detection of B.2 ECVs in early-diverging ferns (e.g *Ophioglossales*) supports infection of the euphyllophyte LCA. In contrast, their restriction in lycophytes to the *Lycopodiaceae* raises the possibility of either ancient vertical transmission or host switching. Thus the most parsimonious hypothesis places the origin of subclade B.2 in the euphyllophyte LCA (∼402 MYA ^38^). Future expanded genomic resources for *Isoetaceae* and *Selaginellaceae* could help resolve this origin, although much of the historical lycophyte biodiversity is extinct ^42,43^.

Taken together, subpatterns of cospeciation are consistent with parallel co-evolution of distinct caulimovirid lineages with their respective host lineages (Figure 6). Still, they are incomplete, likely reflecting historical host switches, ILS, and extinction events. Episomal caulimovirids are largely known from symptomatic plants, and ECVs only record viral lineages that successfully integrated into the germline, leaving an unquantifiable number of interactions undetectable. Recurrent antiviral defenses may have eliminated many lineages independently in different plant groups, while mass extinctions, particularly those at the Late Permian-Triassic (∼250-200 MYA) and Cretaceous–Paleogene (∼66 MYA) ^37,39,42,44^, further eroded signals of cospeciation (Figure 5).

Integrating viral host range with viral phylogeny, we propose four major caulimovirid lineages: Clade A, Clade C, and subclades B.1 and B.2 within Clade B (Figure 6). Our phylogenetic analysis (Figure 2) further suggests that diversification into Clades A, B, and C predates the emergence of subclade B.2, and likely occurred within the euphyllophyte LCA. This scenario supports multiple ancient speciation events coinciding with the Siluro-Devonian expansion of land plants, which may have created new ecological niches for early establishment of the *Caulimoviridae* across tracheophytes.

By expanding both sequence diversity and host range, our study firmly establishes the *Caulimoviridae* as a model system for investigating the origins, diversification, and extinction of plant viruses. As molecular fossils, ECVs uniquely enable the reconstruction of viral macroevolution over timescales spanning hundreds of millions of years, underscoring their crucial role in tracing the deep evolutionary history of plant-virus interactions. Moreover, the evolutionary history of *Caulimoviridae* is deeply intertwined with that of other plant virus families encoding 30k MP proteins, which are unique to plant viruses and essential for cell-to-cell movement through plasmodesmata. 30k MP likely originated from the coat protein gene of a virus infecting an early vascular plant and subsequently spread across disparate viral lineages through horizontal transfer, a transformative event in the emergence of the modern plant virome ^29^. Because many of these MP-encoding lineages predate the *Caulimoviridae*, our work provides a framework for situating the origin and radiation of the modern plant virome within the broader evolutionary history of land plants.

## Methods

### Plant genome datasets

To identify and characterize endogenous caulimovirid sequences (ECVs), we assembled three complementary datasets of plant genomic sequences. The first dataset targeted understudied plant groups to explore novel ECV diversity. It comprised 73 publicly available embryophyte genome assemblies (Supp. Table 1). This dataset included seven bryophyte genomes and 66 genomes from basal tracheophytes: nine lycophytes (three *Isoetaceae*, three *Selaginellaceae*, and three *Lycopodiaceae*), 17 ferns (one *Gleicheniales*, two *Cyatheales*, five *Salviniales*, and nine *Polypodiales*), 27 gymnosperms (seven *Cupressaceae*, two *Taxaceae*, two *Araucariaceae*, eleven *Pinaceae*, two *Gnetaceae*, one *Cycadaceae*, one *Welwitschiaceae*, one *Ginkgoaceae*), three ANA-grade angiosperms (two *Nymphaeaceae,* one *Amborellaceae*), one *Chloranthales*, and nine *Magnoliids* (one *Aristolochiaceae*, two *Magnoliaceae*, two *Calycanthaceae*, four *Lauraceae*). The second dataset was designed to complement the first by including seven angiosperm genomes: *Lindenbergia philippensis*, *Citrus sinensis, Rosa chinensis*, *Ricinus communis*, *Nicotiana sylvestris*, *Vitis vinifera*, and *Glycine max* (Supp. Table 6). A third dataset was compiled, including 14 additional angiosperm genomes representing major lineages: *Dioscorea alata*, *Musa balbisiana*, *Brachypodium distachyon*, *Oryza sativa*, *Papaver somniferum, Nelumbo nucifera*, *Beta vulgaris*, *Vaccinium darrowii*, *Rudbeckia hirta*, *Solanum lycopersicum*, *Capsicum annuum, Arabidopsis lyrata*, *Pyrus communis*, and *Fragaria vesca*, while excluding the *Nicotiana sylvestris* genome (Supp. Table 8).

### Plant transcriptomic datasets

To identify ECVs and *Caulimoviridae* transcripts, we analyzed four publicly available transcriptomic datasets: (1) NCBI Transcriptome Assemblies (https://www.ncbi.nlm.nih.gov/), (2) the “1,000 Plants (1KP)” project by Leebens-Mack (2019; https://sites.google.com/a/ualberta.ca/onekp/) ^24^, (3) a curated set of lycophyte transcriptomes provided by Xia *et al.* (2022) ^26^, and (4) fern transcriptomes compiled by Ali *et al.* (2024; https://conekt.sbs.ntu.edu.sg/species/) ^25^.

### Collection of reference *Caulimoviridae* amino acid RT sequences

A reference library of 106 amino acid reverse transcriptase (aa-RT) sequences homologous to the RT domain of cauliflower mosaic virus (CaMV; GenBank accession number V00141, positions 4,449-5,648) was assembled (Supp. Table 2). It includes 98 aa-RT sequences from *Caulimoviridae* comprising:

i. 16 from episomal viruses, spanning all eleven ICTV-recognized genera ^6^: *Badnavirus* (n=4), *Caulimovirus* (n=1), *Cavemovirus* (n=1), *Dioscovirus* (n=1), *Petuvirus* (n=1), *Rosadnavirus* (n=1), *Ruflodivirus* (n=1), *Solendovirus* (n=1), *Soymovirus* (n=3), *Tungrovirus* (n=1), *Vaccinivirus* (n=1);
ii. 3 from putative caulimovirids: Welwitschia mirabilis virus 1 and 2 (WMV1, WMV2) ^18^ and Pinus nigra virus ^27^;
iii. 79 from ECVs, including: one from the proposed genus Orendovirus, represented by Aegilops tauschii virus ^21^; 11 from the proposed genus Florendovirus ^14^; 56 from the proposed genera Gymnendovirus 1–4 (n=22), Xendovirus (n=2), Yendovirus (n=4), Zendovirus (n=1), Petuvirus-like (n=1), Fernendovirus 1-2 (n=4); 19 from genera *Badnavirus* (n=10), *Caulimovirus* (n=1), *Petuvirus* (n=7), *Soymovirus* (n=1) ^16^; five Beta Endogenous Caulimovirus-like Viruses (BECVs ^17^); six from the proposed genus Wendovirus ^19^.

To root phylogenetic trees, eight outgroup sequences were included: seven from *Metaviridae* and *Retroviridae*, curated in the GyDB database ^22^, and one additional *Metaviridae* RT sequence from *Anthoceros agrestis*.

### Caulifinder libraries

Caulifinder branch A ^20^ is distributed with two core libraries. The “Caulimoviridae_ref_genomes” library, comprising full-length reference caulimovirid genomic sequences, is used for the initial detection of ECVs. The “baits” library, containing ORFs and conserved protein domains from various members of the order *Ortevirales,* is used to filter out false positives ^20,45^. Caulifinder branch B ^20^ is delivered with two distinct libraries. “Caulimo_RT_probes” contains caulimovirid aa-RT sequences used for the initial detection of ECRTs. “Tree_RT_set” includes aa-RT sequences from *Caulimoviridae* and other *Ortervirales*, and serves to filter out false positives and to generate phylogenetic trees of the detected ECRTs.

To improve detection sensitivity and taxonomic resolution, the complete genome sequences of Gymnendovirus 5, 6, 7, and 8, Gnetovirus, and Sequoiavirus ^46^ (Serfraz, personal communication), were integrated into Caulifinder branch A reference genome library. All translated protein sequences ≥ 100 aa in length were incorporated into the “baits” library. The corresponding aa-RT sequences were also added to both the “Caulimo_RT_probes” and “Tree_RT_set_library” of Branch B. To further enhance discrimination between *bona fide* viral sequences and unrelated elements, we supplemented the “baits” library with 545 RT and pol domain sequences from *Metaviridae* and *Retroviridae*, retrieved from the GyDB database ^22^. These augmented resources are hereafter referred to as customized libraries.

### Detection and Clustering of endogenous caulimovirid aa-RT domains (aa-ECRTs)

To identify amino acid sequences corresponding to endogenous caulimovirid RT domains (aa-ECRTs), Caulifinder branch B ^20^ was applied to 73 plant genome assemblies using customized RT libraries. The pipeline was also run on transcriptomic datasets from the 1 KP project and curated lycophyte and fern transcriptomes. In parallel, a tBLASTn search was conducted against the NCBI TSA database using the same custom RT sequences. Due to the database size, this search was restricted to transcriptomic data from five major plant groups: Bryophytes (taxid:3208), Lycophytes (taxid:1521260), Ferns (taxid:241806), Gymnosperms (taxid:1437180), and Angiosperms (taxid:3398). Representative aa-ECRT sequences from these sources were combined with the reference library of 106 RT amino acid sequences.

Pairwise similarity comparisons were performed using BLASTp ^47^, and the results were used to cluster sequences with the Markov Cluster Algorithm (MCL) ^48^ at 62% identity (inflation parameter I = 2). This cut-off was selected to align with genus-level clustering, ensuring clusters approximate recognized caulimovirid genera while resolving distinct OTUs within them. Representative sequences for each cluster were selected based on : (i) a minimum length of 150 amino acids, (ii) presence of the conserved ‘DD’ catalytic motif (aligned to positions 145–146 of the CaMV RT domain) and (iii) good alignment quality.

### Phylogeny of the *Caulimoviridae*

The phylogenetic reconstruction of the *Caulimoviridae* family was based on nucleotide sequences encoding the RT-RH1 domain homologous to that of the cauliflower mosaic virus (CaMV, GenBank accession number V00141, positions 4,449-5,648), the type virus of genus *Caulimovirus* ^6^. To investigate novel OTUs, homologous sequences were recovered from plant genome and transcriptome datasets. Representative sequences were compiled from three sources:

i. Plant genomes: the “sequence retriever” module of Caulifinder branch A ^20^ was run with default parameters on the original 73 embryophyte genomes (Supp. Table 1) plus seven additional angiosperm genomes: *Lindenbergia philippensis*, *Citrus sinensis*, *Rosa chinensis*, *Ricinus communis*, *Nicotiana sylvestris*, *Vitis vinifera*, and *Glycine max* (Supp. Table 6). Consensus sequences were generated when ≥ five ECV copies shared ≥85% nucleotide identity; only those spanning the RT-RH1 domain were retained. Sequences exhibiting <41% BLASTp identity to reference aa-ECRT sequences were excluded, a threshold set to the highest similarity between a caulimovirid RT and that of the *Metaviridae* Ty3, ensuring conservative filtering. Sequences were manually selected by retaining consensus with at least three copies covering the RT-RH1 domain.
ii. Genomic copies: when no or a single consensus could be obtained, unique ECRT loci were extracted using positional information from Caulifinder branch B ^20^ via the “marker miner” module. A 2 kbp flanking region was extracted on both ends to ensure complete domain recovery.
iii. Transcriptomic data: for *Brainea insignis*, for which no complete genome sequence was available, an RT-RH1 domain-containing sequence was retrieved from transcriptomic data (NCBI, accession: GFUE01038349.1).

Sequences were assigned to OTUs by BLASTx (e-value ≤ 1e^-5^) against the 361 aa-RT sequences used to define the OTU with ≥ 62% identity over ≥ 150 residues. Nucleotide sequences < 1,000 bp or producing long branches in preliminary phylogenetic trees were discarded. Up to four representatives were kept per OTU. Alignments used MAFFT v7.511 ^49^ with the G-INS-i strategy and 1,000 iterative refinements. Several alignment trimming strategies were evaluated; however, the untrimmed alignment yielded the most robust and accurate phylogenetic signal. Phylogenetic trees were subsequently inferred using two complementary methods:

i. Maximum Likelihood (ML) with IQ-TREE v2.1.2 ^50^, 1,000 ultrafast bootstrap replicates, best-fit substitution model (GTR+F+I+G4) via ModelFinder ^51^.
ii. Bayesian Inference with MrBayes v3.2.7 ^52–54^, under the GTR+I+Γ model using 2,000,000 generations, sampling every 500. Convergence was assessed using Tracer v1.7.2 ^55^, with 10% burn-in before consensus tree calculations.

### Quantification and classification of nt-ECRTs across plant genomes

To quantify nt-ECRTs across our plant genome dataset, each nt-ECRT identified by Caulifinder branch B ^20^ was queried against the representative aa-ECRT library used to define OTUs, using BLASTx (e–value ≤ 1×10⁻⁵). Assignments to OTUs were made based on each nt-ECRT’s top BLASTx hit, requiring ≥ 65% amino acid identity over ≥ 150 residues (the length of the smallest reference aa-RT domain). We adopted a 65% identity cutoff, more stringent than the 62% threshold used for clustering aa-RTs into OTUs, to minimize misclassification and improve specificity in quantifying nt-ECRT abundance across plant genomes.

### Classification of RT sequences detected in transcriptomic data

The classification of nt-ECRTs recovered from transcriptomic datasets was conducted using BLASTx against the set of aa-RT sequences used to define OTUs.

Each transcript-derived nt-ECRT was assigned to an OTU based on its best BLASTx hit, provided the alignment met a minimum identity threshold of 62% across ≥ 150 amino acids, matching the criteria used during OTU clustering. This consistent framework allowed reliable classification of transcript-derived sequences within the established *Caulimoviridae* diversity landscape.

### Viral genome assembly from ECVs

The WolV1 genome was assembled from ECVs identified in the *W. nobilis* genome, as defined previously ^14^. Briefly, the 5’ and 3’ genomic positions of nt-ECRTs assigned to OTU 19 were extended by 10 kbp, and the sequences of the corresponding loci were extracted. These sequences were compared using an all-against-all BLASTn search to identify high-scoring sequences from different loci that contained identical or near-identical sequences. Fragments of the virus sequence were assembled using VECTOR NTI Advance 10.3.1 (Invitrogen), operated with default settings, except that the maximum clearance values for error rate and maximum gap length were increased to 500 and 200, respectively.

### Phylogeny of *Ortervirales*

To investigate the evolutionary relationships within *Ortervirales*, a representative set of 52 amino acid sequences corresponding to the RT domains of viruses from its five recognized families (*Belpaoviridae*, *Pseudoviridae*, *Retroviridae*, *Metaviridae*, and *Caulimoviridae*) was compiled ^7,22^ (Supp. Table 7).

Sequences were aligned using MAFFT v7.511 ^56^ with the G-INS-i global alignment strategy and up to 1,000 iterations. An ML phylogeny was reconstructed using IQ-TREE v1.6.12 ^50^, with 1,000 bootstrap replicates for branch support. The rtREV+G4 substitution model was selected automatically using the Bayesian Information Criterion (BIC) via ModelFinder ^51^.

### Phylogeny of the 30K movement protein (MP) family

The phylogenetic analysis of the 30K movement protein (MP) family was conducted using amino acid sequences derived from *Caulimoviridae* and homologs from other plant virus families. MP domains from caulimovirid nucleotide sequences were initially identified by annotating protein domains in ECV loci and consensus sequences, and reference genomes using rpsblastn ^47^ with the CDD database (e–value ≤ 1) ^57^. Ambiguous matches were further validated against the Pfam database using HMMER3 ^58,59^. A total of 46 sequences representing the diversity of Clades A, B, and C OTUs were selected and translated, ten of which belonged to Clade C. MP domains retrieved from plant genomic loci were classified following the OTU assignment of their most proximal ECRT within a range of 5 kbp upstream and downstream, following the positions encoding RT and MP domains in ICTV-characterized genomes ^6^.

To contextualize caulimovirid MP diversity within broader viral evolution, an additional 286 MP sequences were sourced from Butkovic *et al.* ^29^, spanning 14 plant virus families: *Alphaflexiviridae*, *Aspiviridae*, *Betaflexiviridae*, *Bromoviridae*, *Botourmiaviridae*, *Fimoviridae*, *Geminiviridae*, *Kitaviridae*, *Mayoviridae*, *Phenuiviridae*, *Rhabdoviridae*, *Secoviridae*, *Tospoviridae,* and *Virgaviridae*.

In total, 332 MP sequences were aligned using MAFFT 7.511 ^56^ with the G-INS-I global alignment strategy and up to 1000 iterations. The alignment was trimmed with trimAl 1.4 using the -gt 0.5 option to remove poorly aligned columns. ML phylogeny was inferred using IQ-TREE 2.1.4 ^50^, with 1000 Sh-aLRT and 1000 ultrafast bootstrap replicates. The LG+F+G4 substitution model was selected based on BIC via ModelFinder ^51^. The resulting tree was midpoint-rooted to facilitate the interpretation of clade relationships.

### Search for Clade C RTs in *Agathis* dammara raw sequencing data

To investigate the potential presence of OTU 19 (Clade C) RT sequences in the genus *Agathis (Araucariaceae)*, we analyzed Illumina whole-genome sequencing data from *Agathis dammara* (SRR15616211), which were initially generated for a mitogenome sequencing project (NCBI bioproject 757934). The dataset comprises 3.4 Gbp, representing approximately 0.12% of the estimated 27 Gbp genome size ^60^. The OTU 19 aa-ECRT rep from *W. nobilis* was used as a query in tBLASTn searches via the NCBI web interface ^23,47^. To maximize detection across the entire RT domain, the probe was segmented into consecutive 50 amino acid fragments, each querying the short-read dataset independently. The top ten hits per fragment were pooled and assembled using CAP3 ^61^, yielding a 529 nt contig, which was translated into a 175 amino acid RT sequence designated as the OTU 19 aa-ECRT rep from *A. dammara* and validated by BLASTx against the curated set of 361 reference RT sequences used in OTU classification.

## Acknowledgements

The authors thank Christophe Plomion (INRAE) for providing access to the *Cupressus sempervirens* genome sequence ahead of publication. The authors thank Anamarija Butkovic for providing access to plant viral 30K MP sequences and Thiery Candresse for critical reading of the manuscript. HV was funded by a joint INRAe-CIRAD PhD fellowship. PYT is funded by the European Regional Development Fund (ERDF, contract REU005756) and the Conseil Régional de La Réunion through the project “Dispositif de partenariat en santé et biodiversité”. URGI is supported by Saclay Plant Sciences-SPS (ANR-17-EUR-0007) and the PlantBioinfoPF platform.

